# A phase oscillator model of cell cycles reveals nuclear density control in a branched fungal network

**DOI:** 10.1101/2025.11.26.690822

**Authors:** Grace A. McLaughlin, Benjamin M. Stormo, Ameya P. Jalihal, Madhav Mani, Timothy C. Elston, Amy S. Gladfelter

## Abstract

Multinucleate cells are widespread in biology, from human skeletal muscle and placenta to many filamentous fungi, yet their basic cell biology remains poorly understood. Maintaining an appropriate nuclear-to-cytoplasmic ratio is essential across cell types for physiological function, and mechanisms of size control have been extensively studied in mononucleate cells. Much less is known about how comparable control is achieved in cells where many nuclei share a common cytoplasm. The filamentous fungus *Ashbya gossypii* forms a branching mycelial network in which individual nuclei divide asynchronously, while the number of nuclei per cell volume (the nuclear density) is tightly controlled. How global regulation of nuclear density coexists with local cell cycle asynchrony remains unclear. We address this by combining a mathematical model of nuclear division in a growing and branching cell with live-cell microscopy. We model nuclei as a dividing population of phase oscillators within a branching cell network and parameterize the model with measurements from *Ashbya* cells. The model demonstrates that asynchrony is required to prevent large density fluctuations that would result from synchronous division, and that introducing a nuclear-density checkpoint to the cell cycles leads to synchrony if it is the only mechanism of density control. We find that coupling branch formation to nuclear density both stabilizes nuclear density and prevents the emergence of synchronous cycles. Our results indicate that asynchronous nuclear cycles together with density-responsive branching maintain a constant nuclear density, revealing a strategy for regulating the nuclear-to-cytoplasmic ratio in large multinucleate cells.

**Significance:** Multinucleate cells appear in diverse biological contexts, from human tissues to filamentous fungi, yet many fundamental aspects of their cell biology are still unclear. Regulating the nuclear-to-cytoplasmic ratio is important across cell types, and little is known about how this is achieved in large, multinucleate fungal cells. Using mathematical modeling and live-cell imaging, we identify how the density of nuclei is controlled in a growing and branching fungal mycelial network. We find that asynchronous nuclear cycles together with density-responsive branching can stabilize the nuclear-to-cytoplasmic ratio within a growing fungal network. This illustrates how large multinucleate cells can control nuclear density even as their morphology becomes increasingly complex.

## Introduction

Multinucleate cells (syncytia) are found across the tree of life and arise in varied ecological and physiological contexts [1]. They play essential roles in human physiology as part of the placenta and skeletal muscle, as well as forming during bone remodeling [2, 3]. Other examples include giant unicellular algae and filamentous fungi, which have critical ecological roles [4, 5]. The formation of multinucleate cells can be either through fusion or nuclear division without cytokinesis. This form of cell organization challenges fundamental principles of cell size control and cell cycle regulation established in mononucleate cells, and presents opportunities for evolutionary innovation by virtue of having multiple, mobile genomes in a shared cytoplasm.

Filamentous fungi are a critical multinucleate cell-type for terrestrial ecosystem function. They grow as networks called mycelia that are composed of filaments termed hyphae, which elongate at their tips and branch to expand the area of the network [6, 7]. Many filamentous fungi do not undergo canonical cell division where nuclear division is followed by cytokinesis, rather individual nuclei within the mycelium progress through the cell cycle and divide without cytoplasmic division. Depending on the species, branches can form either through the hyphal tip forking or emerging laterally off of a hypha [8]. There is significant variability in network topology across fungal species, as ecological context influences how often branches form and the balance of lateral and apical tip branching. Some species have evolved dense, compact networks, while others prioritize foraging by expanding outwards towards new territory [9, 10, 11, 12, 13]. Additionally, even within a given species, the mycelial network of individual fungal cells is highly adaptable, with network structure able to dynamically change in response to external cues such as nutrient availability or physical damage [14, 15, 16]. Furthermore, fungi also adjust their branching patterns for internal functional reasons, such as optimizing cytoplasmic flow and mixing [17]. However, significant gaps remain regarding the mechanisms and timing of branch formation [13].

Large fungal networks can show long-range communication across the cell, even while mounting spatially localized responses to the environment [18, 19]. To coordinate responses, many fungal species display fast cytoplasmic streaming that facilitates long-range nutrient and signal transport throughout the whole cell [20]. Interestingly though, cytoplasmic streaming has also been shown to generate eddies within specific regions of hyphae, which can locally trap nuclei that subsequently become functionally specialized [21]. The positioning of branch points that generate new hyphae is often regulated at the level of the whole cell to maintain average characteristic branch spacing, yet individual hyphae can behave autonomously in response to environmental cues such as nutrient gradients or physical obstacles [22, 23, 24]. Thus, fungal networks balance collective coordination with local autonomy, raising the question of how these opposing features are both achieved within a single cell.

One process where this balance between global coordination and local response is seen is in the process of nuclear division. Notably in the fungus *Ashbya gossypii*, nuclei divide at different times from their spatial neighbors along with showing substantial variation in cell cycle lengths, even while sharing the same cytoplasm [25]. Despite this autonomous behavior of individual nuclei, the number of nuclei per cell volume (nuclear density) is tightly controlled in *Ashbya*. Nuclei are non-randomly positioned throughout the cells, and although local spacing between nuclei has been shown to vary with cell cycle state, the global nuclear density across the entire cell network is tightly regulated [26]. Even in mutants where nuclear positioning is completely disrupted, leading to clustered nuclei and highly variable nuclear density along a hypha, the overall number of nuclei per total cell volume remains similar to wild-type hyphae [27]. This suggests that nuclear density may be sensed across long length scales within these cells.

Nuclear density in a multinucleate cell is analogous to cell size of a mononucleate cell, in that it is a measure of the amount of cytoplasm per nucleus. In mononucleate cells, maintaining a characteristic cell size is critical for proper physiological function, and cell growth is tightly coupled to the cell division cycle to maintain size across the population [28, 29, 30, 31]. This occurs through cell cycle regulation or checkpoints that pause cell cycle progression until the cell reaches an adequate size [32]. Cell volume in *Ashbya*, however, continuously increases as the mycelial network expands and nuclear number increases throughout the cell. Nuclear density is therefore presumably a parameter being controlled in these cells, rather than total volume. Nuclear density has been shown in other fungal species to be relevant in regulating processes such as enzyme production [33]. We hypothesize that, in *Ashbya*, asynchronous cell cycles contribute to nuclear density control by preventing large fluctuations that would arise if many nuclei divided simultaneously. How though are nuclei able to remain asynchronous if they are subject to global nuclear density regulation? To test our hypothesis and answer this question, we developed a mathematical model in which nuclei are represented as a population of dividing phase oscillators within a growing and branching cell. In addition, we acquired time-lapse microscopy movies of nuclei within *Ashbya* mycelia to parameterize and validate the model. Together, these suggest asynchrony and branching morphology are central to maintaining a constant nucleus to cell volume ratio in these large multinucleate cells.

## Results

### Ashbya cells maintain constant nuclear density through time

Single *Ashbya* cells form a large mycelial network that contains many nuclei residing in a continuous cytoplasm (Figure 1A). The hyphae grow exclusively at their tips and occasionally tips split to form two branches (Figure 1B) while nuclei divide throughout the cell network. Previous measurements taken at single time points showed that *Ashbya* cells appeared to maintain similar average nuclear densities, defined as the number of nuclei per cytoplasmic volume [26]. To quantify if indeed nuclear density is stably maintained as cells grow, we imaged fluorescently labeled nuclei through time within individual hyphal networks (Figure 1C, *SI Appendix* Methods). We found that the number of nuclei within the measurable region of the cell was remarkably consistent (Figure 1D, *SI Appendix* Methods). We fit a line to the nuclear density curve for each cell, finding that the resulting slopes were near zero across the population (Figure 1E). Thus, even as a cell grows and changes morphology, the number of nuclei per cell volume is tightly controlled.

**Fig. 1.**
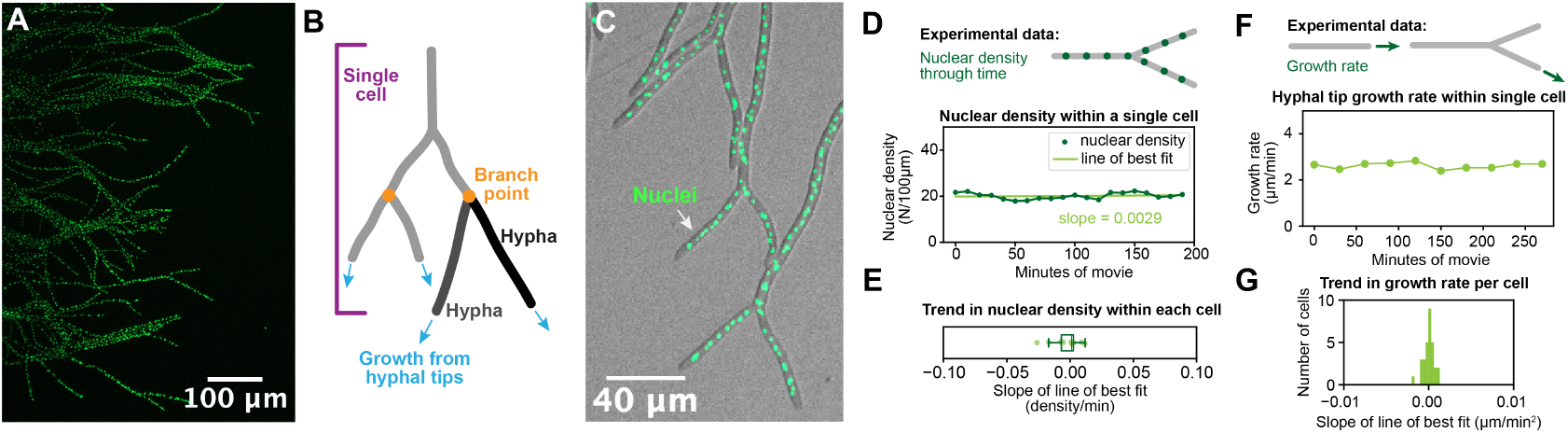
Constant nuclear density motivates mathematical model of nuclear division within a growing cell. (A) Flourescently labeled nuclei within the multinucleate fungus *Ashbya gossypii*. (B) *Ashbya* grows from hyphal tips which periodically fork into new hyphae, which form a large network-like cell structure. (C) Nuclei are imaged within a growing cell. (D) Nuclear density was measured through time over the longest visible region of each branching cell. (E) Nuclear density is held constant through time, as shown by the near-zero slopes of best-fit lines (N = 10 mycelial cell networks, with nuclear density quantified through time across hundreds of micrometers within each). (F) Hyphal tips grow at a steady growth rate, as indicated in (G) by linear fits with slopes near zero when growth rate was measured every 30 minutes (N = 30 cells).

Previous analyses showed that all observed nuclei in these cells can divide, resulting in an exponential increase in total nuclear number [26, 25]. Given the constant nuclear density, a parallel exponential increase in cell volume would be predicted. However, we found the growth rate of hyphal tips through time to be constant (Figure 1F&G, *SI Appendix* Methods). This presents a scaling mismatch, as hyphal volume increases linearly but nuclear number grows exponentially, which would not yield a constant density through time. We hypothesized that branch formation, whereby at some frequency the tips split into two new growth sites, could be sufficient to produce an exponential increase in the cytoplasmic volume needed to keep a constant nuclear density. Are hyphal tip branches sufficient to balance exponential nuclear growth and regulate nuclear density? If so, how frequently must they occur and does this frequency need to be actively coupled to the nuclear division cycle length? To explore whether tip branching provides a possible solution to this scaling mismatch and to evaluate the degree of coordination between growth and nuclear division that leads to a constant nuclear density, we developed a mathematical model of a multi-nucleated fungal mycelium.

### A mathematical model for dividing nuclei within a growing cell

The cell cycle is a biochemical oscillator [34], and our previous work showed that *Ashbya* nuclei progress through this cycle much like those of mononucleate cells [25, 35]. Our focus here is on how nuclear cycle timing interacts with cell morphology to regulate nuclear density, so we model progression through the cell cycle using a phase variable, which captures the timing of division events without requiring explicit molecular detail. Accordingly, we represent a population of nuclei as a growing system of phase oscillators, with cell cycle state denoted by *θ*_*i*_ (*t*) ∈ [0, 2*π*). We assume that each nucleus advances through its cycle at rate 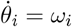, with division occurring when a nucleus reaches 2*π*. At that point we replace the dividing nucleus by two daughter nuclei, both initialized with phase 0, giving a system where the number of oscillators grows through time (Figure 2A).

**Fig. 2.**
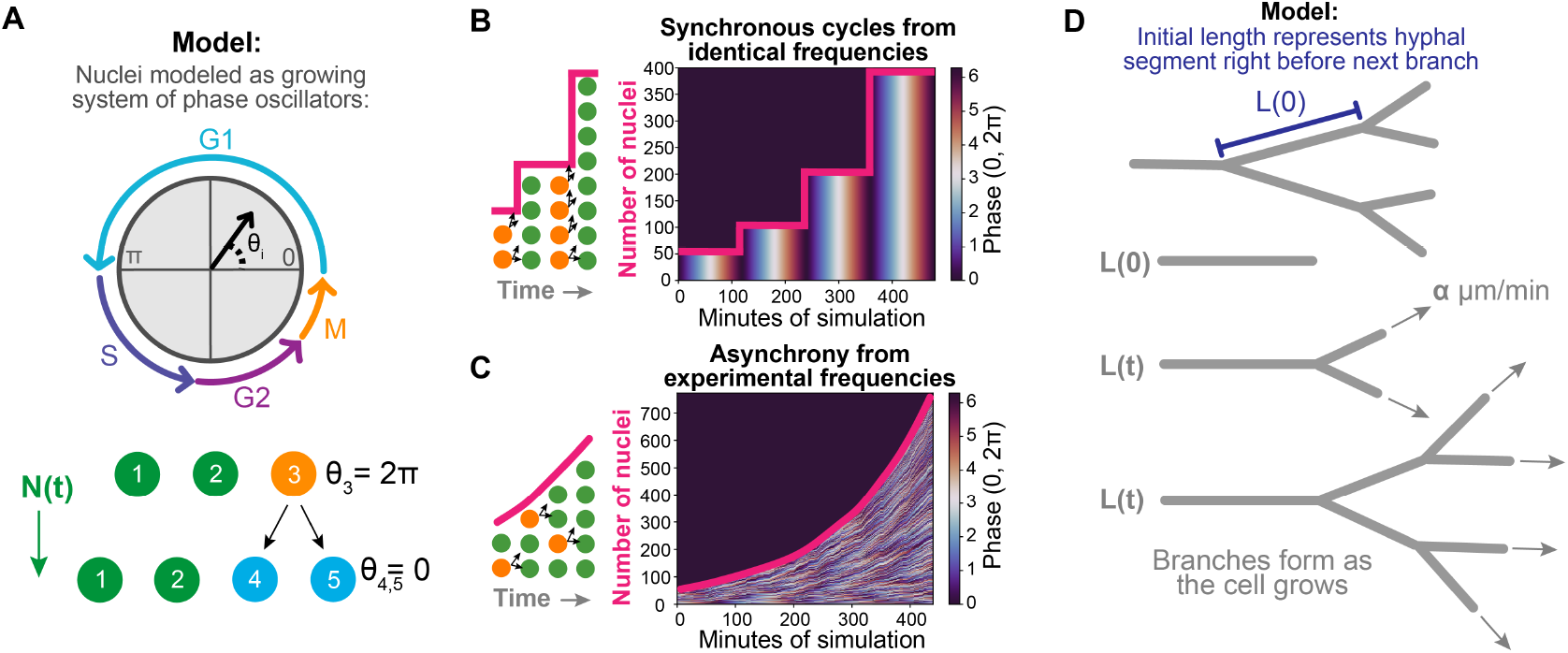
Mathematical model of asynchronous nuclear divisions and branching cell morphology. (A) Nuclei are modeled as a growing system of phase oscillators, with nuclear division occurring at 2*π*. (B) Synchronous divisions occur when each initial phase *θ*_*i*_(0) begins at zero and natural frequencies *ω*_*i*_ are identical and set to 120 minutes. (C) Asynchrony is introduced by assigning each *θ*_*i*_(0) a uniform random phase and drawing each *ω*_*i*_ from a Gaussian distribution with mean and standard deviation of 120 ± 40 minutes. (D) The cell network is modeled as a line that grows from one end and forms branches through time. The initial state represents a hyphal segment immediately preceding the next branch event.

Figure 2B shows the system with identical natural frequencies *ω*_*i*_ where the initial *N*(0) nuclei start at *θ*_*i*_(0) = 0, causing synchronous cycling and sharp jumps in nuclear number at division events. We next assume that the population of nuclei cycle asynchronously, as is the case in *Ashbya*. The mechanisms underlying asynchrony are beyond the scope of this work, therefore we explicitly introduce asynchrony into the model by sampling the phases of the nuclei present at time zero, *θ*_*i*_(0), from a uniform distribution on [0, 2*π*) and drawing each frequency *ω*_*i*_ from a Gaussian distribution whose mean and standard deviation are similar to those of experimentally measured *Ashbya* cell cycle frequencies [26]. This staggers divisions temporally, leading to a smooth, exponential increase in the number of nuclei through time as shown in Figure 2C.

To incorporate cell growth and test the role of branching in stabilizing nuclear density, we next model an *Ashbya* cell as a one-dimensional network that elongates and branches at its tips, mimicking polarized hyphal growth. We define nuclear density as the number of nuclei *N*(*t*) per total length of the cell *L*(*t*). The initial length *L*(0) represents a hyphal segment immediately before formation of the next tip branch (Figure 2D, top). We determined the relevant range of initial lengths *L*(0) by acquiring time-lapse movies of growing cells and measuring the distances between successive branches (*SI Appendix*, Figure S1A). We initialize simulations with the average experimentally measured nuclear density (*SI Appendix*, Figure S1B), adjusting *N*(0) as necessary.

Because we are interested in density control on the scale of the whole mycelium, we do not track positions of individual nuclei. Based on experimental evidence showing long-range sensing of nuclear density and even spacing of nuclei [27, 26], we assumed a uniform nuclear density throughout the modeled cell. As *Ashbya* cells grow, cytoplasmic flow carries nuclei through the network toward the growing tips. We therefore include an influx to *N*(*t*) to account for nuclei entering from connected regions of the cell not explicitly modeled, as this supply also contributes to nuclear numbers in experimental movies (more details in *SI Appendix*, Methods).

The growing ends of our modeled hyphae periodically split into new branches, giving an exponentially growing number of hyphal tips (Figure 2D, bottom). Total cell length increases through time at rate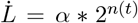, with *α* denoting the growth rate of individual hyphal tips determined from the mean value obtained from experimental measurements (*SI Appendix*, Figure S1C). 2^*n*(*t*)^ gives the number of growing tips, where each time a branching event occurs, *n*(*t*) increases by one. The time intervals between these branching events are treated as a random variable *T*, whose distribution is determined from experimental data (Figure 3A). Model variables are summarized in Table 1 and model parameters in Table 2.

**Table 1.**
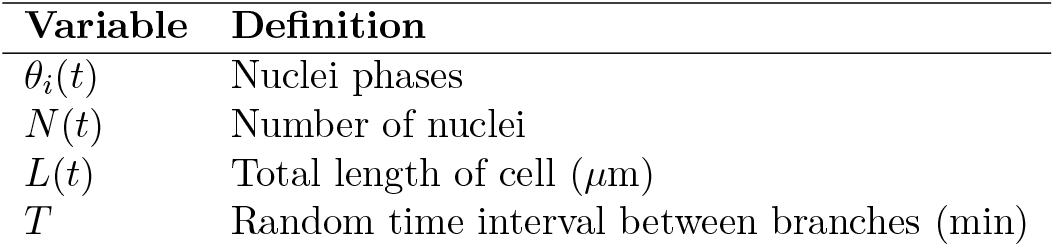
Model variables used in simulations.

| Variable | Definition |
| --- | --- |
| $\theta_i(t)$ | Nuclei phases |
| $N(t)$ | Number of nuclei |
| $L(t)$ | Total length of cell ($\mu\text{m}$) |
| $T$ | Random time interval between branches (min) |

**Table 2.** Model parameters used in simulations.

| Parameter | Definition | Value, range, or distribution |
| --- | --- | --- |
| $\theta_i(0)$ | Initial oscillator phases | Drawn from $\mathcal{U}(0, 2\pi)$ |
| $N(0)/L(0)$ | Initial nuclear density (N/100$\mu\text{m}$) | 20 |
| $L(0)$ | Initial cell length ($\mu\text{m}$) | Ranging from 200–500 |
| $N(0)$ | Initial number of nuclei | Ranging from 40–100 |
| $\alpha$ | Cell growth rate ($\mu\text{m}/\text{min}$) | 2 |
| $\omega_i$ | Cell cycle natural frequencies (1/min) | Drawn from $\frac{2\pi}{\mathcal{N}(\mu_\omega, \sigma_\omega)}$ |
| $\mu_\omega, \sigma_\omega$ | Mean, SD of cell cycle lengths (min) | 120, 40 |
| $\mu_T, \sigma_T$ | Mean, SD of branch intervals (min) | 120, 40 |

**Fig. 3.**
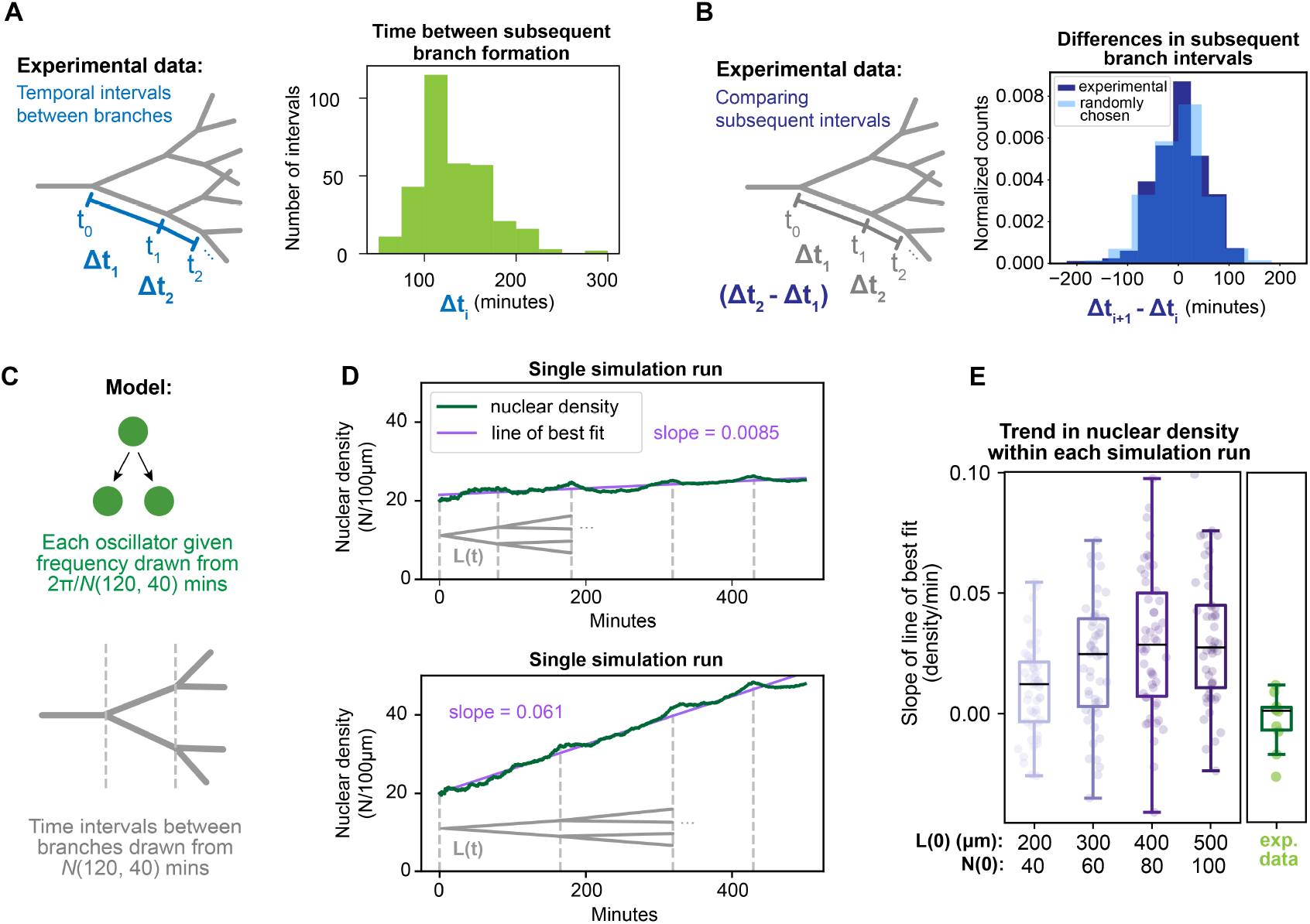
Cell cycle lengths matching branching times is not sufficient for constant nuclear density. (A) Time intervals between branching events were measured in *Ashbya* cells (N=326 temporal intervals across 93 cells). (B) Dark blue shows the difference between the timing of subsequent branches within individual cells (N=233 time differences). This is compared to the difference between values randomly sampled from the distribution in A (N=10,000). (C) In the model, nuclear cycle lengths and branch intervals are drawn from identical Gaussian distributions. (D) Example simulation runs with *L*(0) = 400. (E) Slopes of linear best fit to nuclear density curves, 50 simulation runs per *L*(0). Each simulation was run for 500 minutes. Experimental data from Figure 1E shown to the right.

### Measured branch timing is similar to cell cycle timing

If branching cell geometry is involved in controlling nuclear density, then the timing of branch formation may be a critical part of this coordination. To ensure the number of branching events *n*(*t*) evolves in our model in a manner consistent with experimental observations, we measured the temporal intervals between tip branch formation in *Ashbya* (*SI Appendix*, Methods). These intervals were measured in mature cells that are no longer producing lateral branches. Interestingly, we found that the branch-timing distribution with mean and standard deviation of 130 ± 39 minutes is strikingly similar to the cell cycle length distribution in *Ashbya* cells of 118 ± 38 minutes (Figure 3A) [26]. The similarity in both mean and variability suggests that branch formation and nuclear division operate on matching timescales. Moreover, branch timing within individual cells is highly variable, and we did not find correlation between subsequent branch intervals in the same cell (Figure 3B).

Because maintaining constant nuclear density requires exponential increases in nuclear number to be balanced by comparable increases in total cell volume, these observations motivated a simplified model in which the two processes share the same underlying timing distributions. We therefore first set the model so that both the cell cycle lengths and the times between branching events, *T*, are drawn independently from a common Gaussian distribution of 120 ± 40 (SD) minutes. This allows us to directly test the fundamental question of whether matching the timing distributions of nuclear division and branch formation is sufficient to stabilize nuclear density.

### Active coordination is required for nuclear density control

To determine whether the model with matched timing distributions of the cell cycle and branching can maintain constant nuclear density, we performed simulations (Figure 3C). A small subset of the model simulations produced a nearly constant nuclear density (Figure 3D, top). In general, however, density is not maintained by this model (Figure 3D, bottom). Therefore, treating cell cycle progression and branching as independent random processes with identical distributions is insufficient for robust nuclear density control (Figure 3E). We note that increasing the initial length (and therefore the number of nuclei) amplifies the trend of increasing density, as the system begins further along the exponential growth curve of nuclear number. Therefore, simply matching the timing of cell cycle progression and branch formation is not sufficient to maintain constant nuclear density across all cells. An active mechanism is required to keep the exponential increase in nuclear number aligned with the exponential increase in growing tips, even if their relative rates of increase match.

We do not know the extent to which variability in cell cycle lengths in *Ashbya* comes from intrinsic natural frequency differences or through factors in the shared cytosol such as noisy concentration gradients of cell cycle regulators. However, if we assume identical oscillators and introduce Gaussian white noise to generate variability, the model produces results similar to those presented here (*SI Appendix*, Figure S2A&B). Furthermore, our model assumes that each generation of new branches forms simultaneously. In reality, individual branches form at different times, but this variability cannot be fully quantified in the microscopy movies due to overlapping hyphae during growth. If we assume that the timing distribution between every branch event follows the same Gaussian distribution in Table 2, we similarly find that nuclear density is not well controlled (*SI Appendix*, Figure S2C&D). These results indicate that simply having divisions and branching occur on similar timescales is insufficient for density control and instead active regulatory mechanism(s) are required to maintain constant nuclear density.

### A cell cycle checkpoint leads to synchrony

In many systems the cell cycle is connected to other processes such as growth through a series of checkpoints. These monitor key events related to division and police progression through the cell cycle so that it proceeds only when necessary conditions are satisfied. In budding yeast, a close evolutionary relative of *Ashbya*, monitoring during the growth phase G1 prevents entry into S phase until the cell has reached a sufficient size [36, 32, 37]. Because *Ashbya* cells do not divide but rather continuously increase in volume, we hypothesized that these cells sense nuclear density instead of cell volume. We therefore added a checkpoint to our phase oscillator model that pauses cell cycle progression if nuclear density exceeds a threshold, which is analogous to a cell being too small (Figure 4A). We define G1 as *θ*_*i*_(*t*) ∈ [0, *π*), as G1 spans approximately half of the *Ashbya* cell cycle [35]. We set the density threshold to 23 nuclei/100*µ*m, approximately three standard deviations above the experimental mean value of 20 nuclei/100*µ*m, based on the average standard deviation of nuclear density fluctuations measured through time. As each *θ*_*i*_(*t*) reaches *π*, if *N*(*t*)*/L*(*t*) exceeds the threshold, that nucleus is temporarily paused at *π*. When nuclear density drops below the threshold, the cell cycle resumes with frequency *ω*_*i*_, as defined in Table 2. We also assume that escape from arrest can occur stochastically before nuclear density falls below the threshold, based on the observation that budding yeast exhibit imperfect size regulation [32]. Premature escape from G1 is modeled as a Poisson process with rate *λ* (see *SI Appendix*, Methods for details). Temporal intervals between branching events are randomly sampled from the experimental distribution as described previously. We use our extended model to determine if addition of a cell cycle checkpoint leads to nuclear density control.

**Fig. 4.**
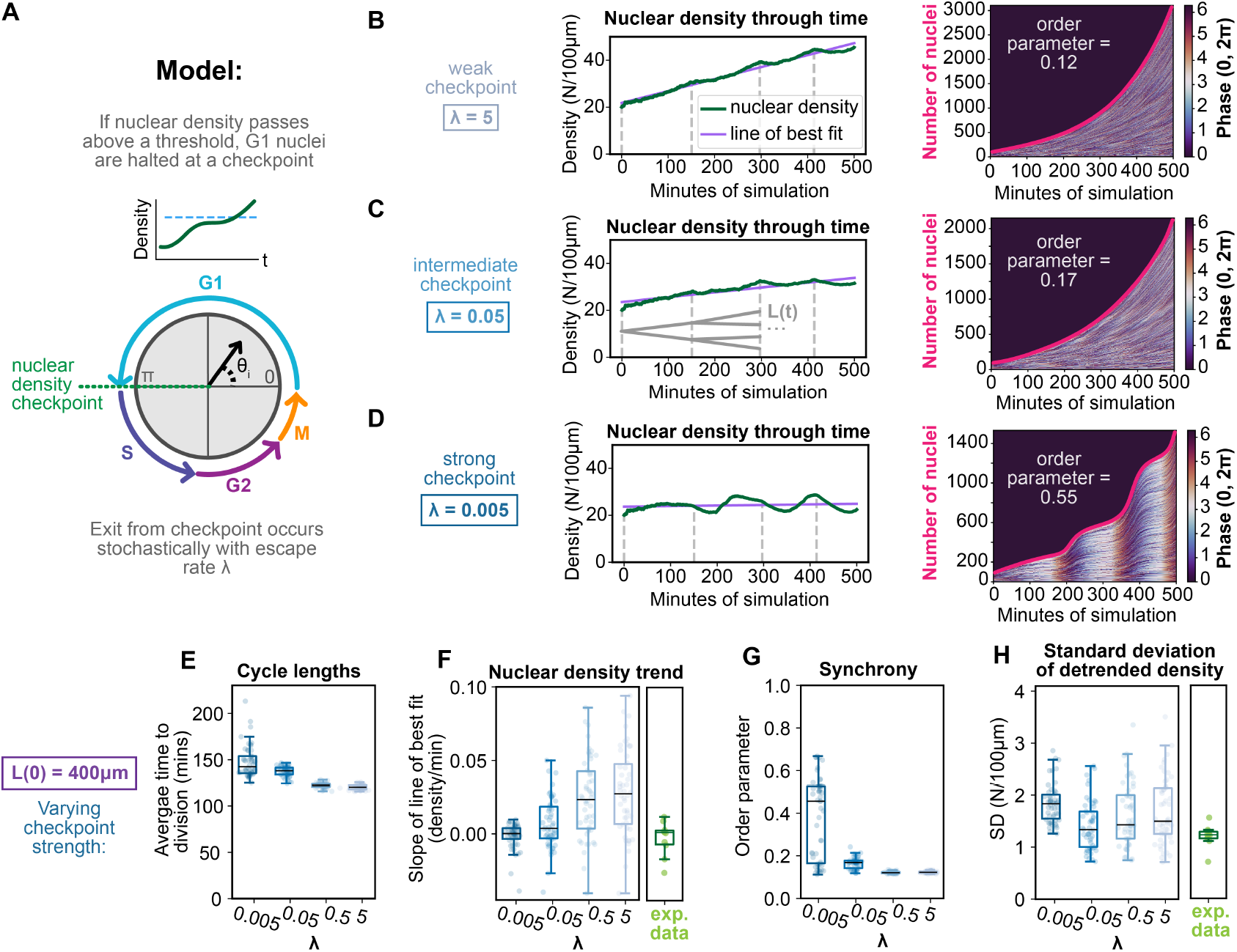
Cell cycle checkpoint synchronizes nuclear phases. (A) A nuclear density checkpoint is added to the model such that nuclei pause oscillation at *θ*_*i*_ = *π* if nuclear density is above a threshold value. Stochastic exit from the checkpoint while density is above the threshold occurs at an average rate *λ. L*(0) is fixed at 400*µm*. (B-D) Example simulations showing the effect of decreasing *λ*. (E) Average cycle length per simulation (N=50 per *λ* value). Lowering *λ* lengthens cycles as nuclei are held at the checkpoint longer on average. (F) Slopes of linear best fit to density. Decreasing *λ* improves this metric of density control. (G) Lower *λ* values lead to synchronous cycles, quantified by an average order parameter per simulation (more details in *SI Appendix*). (H) Synchronous divisions increase the standard deviation of density values around the best fit line.

A high G1 escape rate (*λ* = 5) causes nuclei to pass the checkpoint rapidly, functionally eliminating it and leading to poor control of nuclear density (Figure 4B). Decreasing *λ* to 0.05 strengthens the checkpoint, improving density control (Figure 4C). However, as we decrease *λ* further to 0.005, the slope of the nuclear density trend approaches zero (Figure 4D, left), but the phases begin to synchronize as oscillators accumulate at the checkpoint (Figure 4D, right). Synchrony leads to nuclear divisions being temporally clustered, resulting in sharp increases in nuclear density. To quantify synchronization between nuclei, we calculate the order parameter 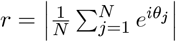 at each time step [38], and then take the time average to obtain a single value per simulation (details in *SI Appendix*, Methods).

Over many simulation runs we found that lower *λ* values slow cell cycle lengths (Figure 4E) which prevents nuclear density from trending upwards (Figure 4F). This, however, synchronizes nuclear cycles (Figure 4G). Synchrony produces larger fluctuations in density compared to the experimental data (Figure 4H). This result suggests that *Ashbya* nuclei must divide asynchronously to minimize density fluctuations, consistent with experimental observations that neighboring nuclei are often in different stages of the cell cycle [25].

As *λ* is decreased, the model produces cycle lengths longer than what the natural frequencies would give rise to alone. We next shift the natural frequencies to attempt to match the experimental average cell cycle cycle length. For the intermediate checkpoint (*λ* = 0.05), decreasing the mean natural cycle length from 120 to 100 minutes yields a resultant cycle length comparable to the experimental value, but at the cost of diminished density control (*SI Appendix*, Figure S4A-C). For a stronger checkpoint (*λ* = 0.005), regardless of the natural cycle length, density is well controlled, but the resulting cycle lengths remain higher than the experimental value (*SI Appendix*, Figure S4D-F). We find similar results if each branch event occurs independently, rather than new generations of branches forming simultaneously (*SI Appendix*, Figure S4G-L).

We have shown that strengthening the cell cycle checkpoint to the point that it prevents increases in nuclear density causes nuclei to become more synchronized, which in turn produces large temporal fluctuations around the mean nuclear density. Therefore, our model is still missing a mechanism that stabilizes nuclear density in a way that matches the experimental observations. Per simulation, there are hundreds of nuclear division events but fewer than ten branching events. As a result, if both nuclear cycles and branch times are drawn randomly, each simulation samples the distribution of nuclear cycle times well, whereas branch timing is determined by only a handful of random values. Consequently, the few randomly chosen branch intervals disproportionately influence the outcome of each simulation run. This sensitivity, together with the inability of the checkpoint model to reproduce the experimental data, suggests that branch formation is unlikely to be purely stochastic, leading us to hypothesize that it is instead regulated by nuclear density.

### Density-dependent branch formation maintains constant nuclear density

To test the hypothesis that timing of branch formation is coupled to nuclear density, we update the model to have branch formation triggered when nuclear density exceeds the threshold used in the checkpoint model (Figure 5A). We assume that branch formation does not occur instantaneously but rather follows a Poisson process with rate *β* once nuclear density exceeds the critical threshold. Furthermore, we assume a minimum time between branch events based on the lower end of the experimental distribution of branch times (more details in *SI Appendix*, Methods). When *β* = 1, branches form rapidly, and we found significant improvement in nuclear density control. Branch formation now does the bulk of the work controlling density, leading to less reliance on the cell cycle checkpoint and reduced slowing of cycle lengths (Figure 5B). Across checkpoint strengths (*λ*), nuclear density is well controlled (Figure 5C) and asynchrony is maintained (Figure 5D).

**Fig. 5.**
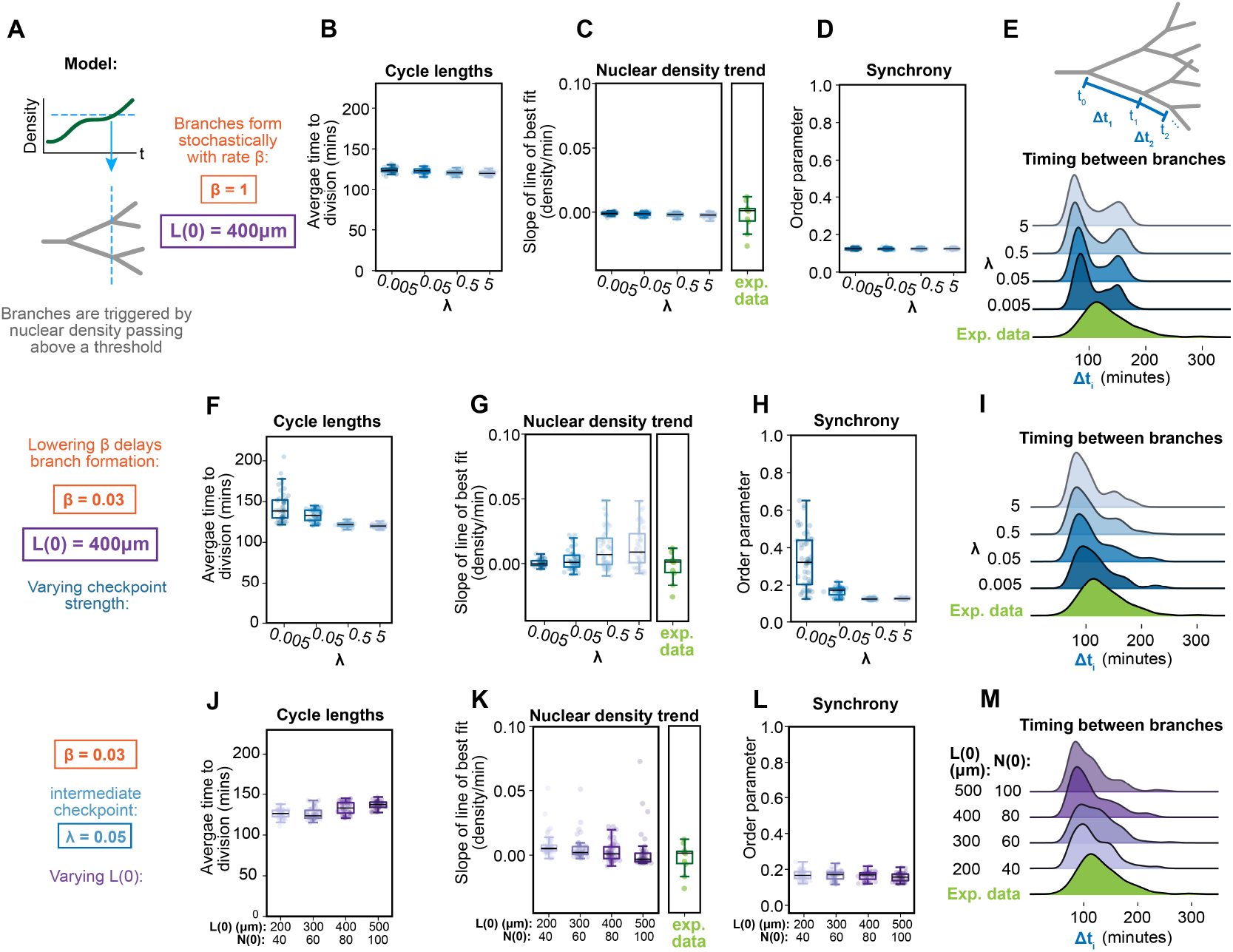
Regulated branch formation improves nuclear density control. (A) In the model, sets of new branches now form at rate *β* when nuclear density exceeds a threshold value. We initially fix *L*(0) = 400*µm*. (B) Branches form rapidly when *β* = 1, leading to less reliance on the cell cycle checkpoint for density control. Average cycle lengths per simulation remain similar across checkpoint strengths (N=50 runs per *λ* value), (C) slopes of the linear best fit to density remain close to zero, and (D) asynchrony is maintained, quantified by an average order parameter per simulation run. (E) Distributions of time intervals between branch formation for each *λ* value are visualized as kernel density estimates and compared to the experimental distribution from Figure 3A. (F-I) The same metrics as panels B-E are shown for *β* = 0.03. This delay to branch formation leads to better overall matching to experimental data, so long as the checkpoint strength is intermediate (*λ* = 0.05). (J-M) We fix *β* = 0.03 and vary *L*(0) across values in Table 2, finding nuclear density is well controlled across initial conditions.

To assess whether the model could produce a distribution of branch intervals similar to the experimental distribution, we examined the distribution of branch timing (Figure 5E). We found that the model produced a left shifted, bimodal distribution of branch times due to a series of branches forming rapidly at the onset of each simulation. We do not observe this in the experimental data. Lowering *β* reveals a trade off: as we increase the delay in branch formation, the distribution of branch intervals more closely matches the experimental data, but density control is worsened (a range of *β* values is shown in *SI Appendix*, Figure S5). The best balance is produced with *β* = 0.03. We found that as long as the cell cycle checkpoint is of intermediate strength (*λ* = 0.05), nuclear density is well controlled, the system maintains asynchrony, and the branch timing distribution with a mean and SD of 113 ± 37 minutes is similar to the experimental distribution (Figure 5F-I). If the checkpoint is too strong (*λ* = 0.005) then the model produces synchronized divisions, suggesting that some stochasticity in checkpoint escape is necessary. If the checkpoint is weak (*λ* = 5) then nuclear density is not well controlled without branches forming faster than is experimentally observed, suggesting that both a cell cycle checkpoint and density-dependent branch formation are required for proper nuclear density control.

Finally, we test whether the model is robust across different initial cell lengths. We fix *λ* to 0.05 and vary *L*(0) and *N*(0) across the ranges in Table 2. As shown in Figure 5J-M, we find similar results across initial conditions, indicating that our model robustly reproduces the experimental data. However, we note that the degree to which the cell cycle checkpoint acts to control density varies with *N*(0), as a larger initial number of nuclei leads to faster exponential growth in nuclear population. This leads to increasing cycle lengths with increasing *N*(0). Furthermore, no single shift in the natural frequencies produces cycle lengths that match the experimentally measured value across all initial conditions (*SI Appendix*, Figure S6). We note that the published average cycle length was measured in shorter hyphae, and therefore predict that the average duration of the cell cycle in *Ashbya* varies through time, such that longer hyphae with more nuclei have longer cycles to prevent increases in nuclear density.

We have shown that coupling tip branch production to nuclear density results in significant improvements to nuclear density control. Furthermore, we found that active coordination in both timing of new branch formation and in cell cycle progression is required to control nuclear density. For the model to capture the experimental data, a balance between these two mechanisms is required. A degree of randomness is also required in both mechanisms: nuclei occasionally escape the checkpoint at high density, and branches form stochastically after a delayed response to density surpassing a threshold. A prediction of the model is that nuclear density is highest preceding new branch formation. We next reanalyze our experimental data to look for evidence supporting this prediction.

### Nuclear density is highest preceding branch formation

To determine if nuclear density peaks preceding branch formation in *Ashbya*, we focused on individual hyphae and measured the number of nuclei within 50 micrometers of the hyphal tips before and after branch formation (Figure 6A). In all hyphae, nuclear density was higher before the branch compared with after (Figure 6B). This indicates that although global density is kept remarkably constant within a growing mycelium, branch formation leads to local fluctuations in nuclear density near hyphal tips. There is, however, variability in the nuclear density preceding a branch (Figure 6C), possibly arising from stochasticity in how rapidly branches form in response to rising nuclear density.

**Fig. 6.**
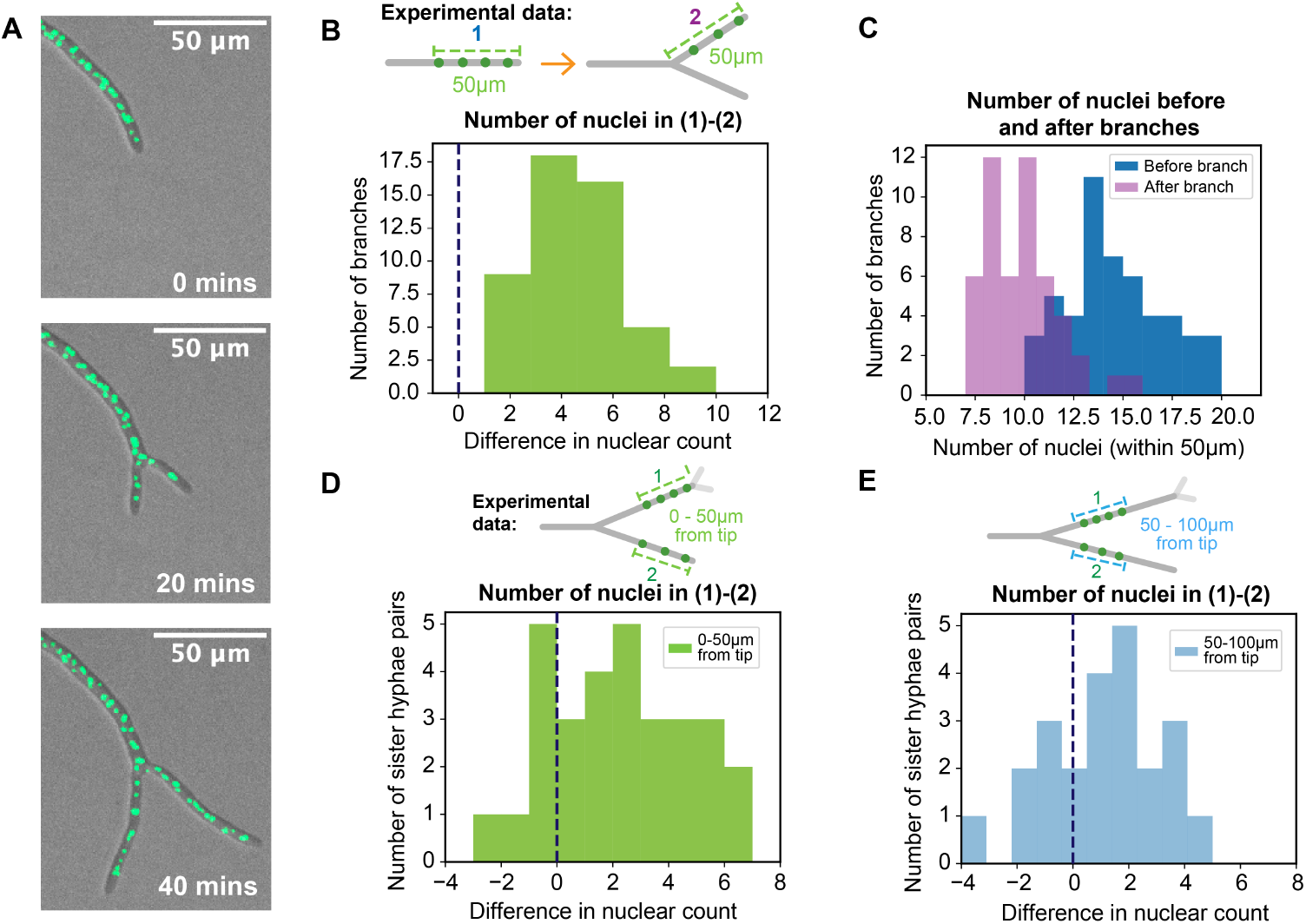
Branch formation locally lowers experimental nuclear density, while rising nuclear density precedes new branches. (A) Nuclei within *Ashbya* cells before and after formation of a new branch. (B) Difference between the number of nuclei before and after branch formation within a 50 *µ*m region of the hyphal tip. In all cells (N= 50) the difference was greater than zero, demonstrating that branching locally drops nuclear density. (C) The number of nuclei within every cell prior to branching (dark blue) compared with after branching (pink). (D) We next focus on both sides of branches within individual cells, still counting nuclei within 50 *µ*m of the hyphal tip. In the diagram, (1) indicates the hyphal segment that undergoes branching first. At the time point prior to this segment branching, we subtract the number of nuclei in the other hyphal branch, labeled as (2) from the number within segment (1). In the majority of pairs (20 out of N=30 pairs), the nuclear count is higher in the hyphal segment that branches first. (E) The number of nuclei also is higher on average in the first branching hypha when measured in a 50 *µ*m region farther back from the tip.

We next sought an internal comparison for further evidence that branching directly responds to rising nuclear density. We looked within individual cells, and focused on instances where after a branch, the next two branches did not form simultaneously. We found that in the majority of cells, nuclear density is higher in the hypha that branches first (Figure 6D). This is most pronounced within 50 micrometers of the hyphal tip, but we still found a tendency toward higher density in the early branching hypha when we compared segments more distal to the hyphal tip (Figure 6E). Taken together, these data suggest that branches function to locally decrease nuclear density, and that nuclear density is coupled to branch formation. The exact length scale over which nuclear density is sensed and the specific molecular signals involved remain open and intriguing questions.

## Discussion

Our goal was to understand how nuclear density is controlled in the mycelium of a multinucleated fungus. To approach this, we combined a mathematical model of dividing nuclei within a branching cell network with live-cell microscopy of growing *Ashbya* cells. Nuclear density control requires that nuclei do not synchronize their division cycles, because this would produce large variations in density as nuclei simultaneously divide. Therefore, we assumed nuclei progress asynchronously through their cycles. The model showed that tip branching generates the exponential increase in cell volume needed to accommodate dividing nuclei. However, branching alone at the experimentally measured frequency could not robustly maintain stable density. Reproducing the experimental data required coupling both cell cycle progression and branching to nuclear density. We found that including a density-dependent cell cycle checkpoint prevented nuclear density increases, but also synchronized the nuclear division cycles, which led to large nuclear density fluctuations. When branching was also regulated by nuclear density, the strength of the cell cycle checkpoint could be diminished such that nuclei remained asynchronous and density was stabilized. Inclusion of this weakened cell cycle checkpoint was necessary for the model to match the experimentally observed distribution of branch time intervals. Consistent with these predictions, our experiments showed that branching reduces nuclear density and that local density increases precede branch formation.

In budding yeast, cell size is regulated largely by a checkpoint at the G1/S transition, where smaller cells extend their time in G1 to allow additional growth [32]. Given the genetic similarity between *Ashbya* and budding yeast [36], it is reasonable to expect that a similar cell cycle checkpoint mechanism operates in *Ashbya*. However, controlling nuclear density within a mycelial network adds complexity, as exponential increases in nuclear number must now be balanced within a single continuous cell. Furthermore, the *Ashbya* cell shape is more expansive than budding yeast, as the mycelial network gains an exponentially increasing number of tip branches over time. Our modeling suggests that *Ashbya* requires more than cell cycle regulation to control nuclear density, with timing of tip branch formation playing a critical role.

As shown in Figure 1D&E, we found that across hundreds of micrometers within an *Ashbya* cell network, nuclear density remains remarkably constant over time. Interestingly though, when we examined density near hyphal tips, we found that tip branch formation led to significant local decreases in nuclear density (Figure 6B). New tip branches locally drop nuclear density by doubling the cytoplasmic volume that nuclei can move into as the hyphae grow, while globally across a mycelium branching serves to stabilize nuclear density by offsetting the exponential growth of nuclear number. In our model we made the simplifying assumption of uniform nuclear density across the cell network, which allowed us to investigate the interplay between branch formation and nuclear density control without introducing the complexity of nuclear movement and positioning. However, as shown in Figure 6D&E, differences in nuclear density do occur between hyphae within the same cell network, suggesting that there is a finite length scale over which density is sensed, possibly limited by diffusion. The ability for individual hyphal tips to autonomously respond to local nuclear density changes may allow for more optimal buffering of global nuclear density fluctuations. An intriguing open question is whether the length scale over which density is sensed for tip branching is similar to that used by nuclei to regulate their cell cycle progression.

Filamentous fungi regulate nuclear division and hyphal growth in various ways, and even within a species these strategies can change over time [13]. *Ashbya* shifts from an early slower growth phase with lateral branching to a mature phase with rapid extension and tip branching (the growth stage we focused on here). How nuclear density is maintained during the lateral branching phase and how *Ashbya* cells transition their nuclear density control mechanisms across growth phases remain open questions. Across species, *Neurospora crassa* also shows asynchronous nuclear cycles and both lateral and tip branching, whereas *Aspergillus nidulans* displays synchronous waves of nuclear divisions and lateral branches that arise only behind septa, which create compartments no longer continuous with the growing hyphal tip region [39]. Furthermore, branch formation has been shown to be coupled to the cell cycle within *Aspergillus nidulans* such that higher nuclear density within a compartment leads to more branches [40, 41]. Furthermore, the total number of nuclei has been shown to scale with cell volume across multiple *Aspergillus* species [33]. These examples hint that hyphal morphology programs and levels of nuclear synchrony may co-evolve, motivating comparative studies across fungal species and developmental stages.

Our modeling results generate several experimentally testable hypotheses about the interplay between nuclear density, cell cycle progression, and branching in a growing fungal network. We predict that cell cycle asynchrony keeps fluctuations in nuclear density low. Published *Ashbya* mutants with more synchronous nuclei show increased variability in distances between neighboring nuclei, which could translate into greater variability in overall nuclear density [42]. Thus, measuring nuclear density through time in these more synchronous strains would be an informative direction for future work. The model also predicts that proper tip branch formation is critical for density control. Thus, we expect mutants with defective branching to show larger fluctuations in nuclear density over time [43]. We further predict that nuclear cycle timing may adapt as cells grow. For example, in longer hyphae with more nuclei, slower cell cycles may be necessary to maintain nuclear density. It would therefore be informative to measure nuclear cycle durations across time within individual growing hyphae to test whether cell geometry feeds back on the cell cycle, although doing so in fast growing, mature hyphae remains technically challenging due to rapid nuclear movement and tip extension. It would also be interesting to determine how environmental conditions, such as nutrient depletion or temperature, affect nuclear density control. Proper nutrient levels may be required for *Ashbya* nuclei to pass through cell cycle checkpoints, in which case nutrient limitation would be expected to slow nuclear cycles. Finally, the molecular basis of nuclear density sensing in *Ashbya* remains unknown. Identifying the signals and pathways that couple nuclear density to both cell cycle progression and branching will be an important direction for future work.

## Materials and Methods

To visualize nuclei in live cells, we integrated a construct containing mNeon fused to the C-terminus of the *Ashbya* H4 gene. Nuclear density fluctuations were measured at high spatial resolution on a Nikon Ti-Eclipse microscope with a Yokogawa CSU-W1 spinning disk. For comparative density measurements within individual cells, we used a Zeiss Axio Zoom widefield fluorescence microscope to capture a larger field of view. The Zeiss Axio Zoom microscope was also used to measure hyphal growth rates and tip branch intervals. Full experimental details, including sample preparation, as well as supplemental information about the mathematical model are provided in the *SI Appendix*. All code will be made available on GitHub upon final publication.

## Acknowledgments

This work was funded by NIH GM R35 GM118096, NIH R35GM127145, and NIGMS 1F32GM147989.

