## Supplemental Appendix for "A phase oscillator model of cell cycles reveals nuclear density control in a branched fungal network"

### Experimental Materials and Methods:

**Constructing fluorescent nuclear strain.** To visualize nuclei live in *Ashbya*, we integrated a copy of the *Ashbya* H4 histone fused in-frame at its C-terminal end to *Ashbya* codon optimized mNeon. Briefly a pRS416 vector containing homology to a safe harbor location on chromosome VI was digested with BstBI (NEB R0519) and NdeI (NEB R0111) generating a 708bp band. The yeast *HIS3* promoter, the Ashbya H4 gene, mNeon, and the yeast *ADH1* terminator were amplified by PCR. All five fragments were run on an agarose gel to confirm size and purity. The five bands were gel extracted according to the manufactures instructions (QiaQuick Gel Extration kit, Qiagen, 28704). All five fragments were mixed in equal molar amounts and assembled by Gibson (NEB, Nebuilder HiFi DNA assembly kit, E2621). Gibson product was transformed into *E. coli* according to the manufactures instructions (NEB 5-alpha competent, C2988) and grown under selection. Individual colonies were picked, grown under selection in liquid media and DNA was extracted using a miniprep kit according to the instructions (thermofisher, GeneJet Plasmid Miniprep, K0502). Correct sequence was confirmed by whole plasmid long read sequencing (Azenta Life Sciences).

To prepare linear DNA for integration PCR was performed on the plasmid using primers that generate a 4160bp fragment containing: >200bp of homology at either end to a safe harbor on the chromosome VI of Ashbya, a KanMX antibiotic resistance gene, and the *Ashbya H4* gene fused to mNeon. The PCR product was checked for purity by agarose gel. Any parent plasmid was removed by digestion with Dpn1 (NEB, R0176). This fragment was transformed into *Ashbya* by electroporation as previously described [1]. Transformants were verified by PCR and Sanger sequencing (Azenta Life Sciences, South Plainfield, NJ, USA).

**Measuring nuclear density fluctuations in growing cells.** High spatial resolution was required to quantify nuclear density changes through time within individual cells, as *Ashbya* nuclei frequently come spatially close to one another. Therefore, for the data in Fig. 1D&E of the main text we imaged nuclei within growing *Ashbya* cells on a spinning disk fluorescent microscope. The H4-mNeon *Ashbya* cells were grown from 10 $\mu$L of spores in 125mL baffled flasks under G418 selection (200$\mu$g/mL) in 10mL *Ashbya* full medium (AFM) shaking at 110rpm for 19 hours at 25 degrees C. Cultures were spun for 3 minutes at 300rpm in 15mL conical tubes, washed with low flourescence media (LFM), and mounted on a gel pad composed of LFM and 1% agarose. The edges of the coverslip were sealed with valap and allowed to rest for 3 hours at room temperature. Slides were them imaged on a Nikon Ti-Eclipse stand with a Yokogawa CSU-W1 spinning disk. Nuclear imaging was performed using a 40x Nikon oil-immersion objective and a 488 nm laser. At each time point, three z-slices were acquired at 1.8 $\mu$m intervals. We then quantified nuclear density fluctuations within individual *Ashbya* mycelial networks. At 10 min intervals, we counted the total number of nuclei per total cell length. The measurable cell length varied over time as cells grew and branched, and measurements ranged from 80 to 400 $\mu$m. When a tip branch occurred, length measurements included both segments. Nuclear density was measured for a minimum of 120 mins per cell network.

**Nuclear density comparisons within individual cells.** For nuclear density comparisons within individual *Ashbya* cell networks in Fig. 6 of the main text, we used an imaging modality with a wider field of view to capture a larger number of cells and to visualize larger portions of individual cell networks. Although this reduced spatial resolution, it was sufficient for our comparative measurements within individual cells. H4-mNeon *Ashbya* cells were mounted on gel pads as described in the previous section, except that a gas-permeable polymer coverslip was used (Ibidi brand). The histone-tagged *Ashbya* strain was imaged on a Zeiss Axio Zoom widefield fluorescence microscope using 488 nm excitation light, with images collected every 10 minutes. For measurements of nuclei near the hyphal tip in Fig. 6, we used the position of the leading nucleus as the starting point and counted the number of nuclei within 50$\mu$m of it.

**Measuring hyphal growth rate and tip branching intervals.** We next measured hyphal tip growth rates and the intervals between tip branches. 20 $\mu$L of spores from the previously described H4-mNeon *Ashbya* strain were placed in the center of AFM plates and allowed to grow for a minimum of 24h at 25 degrees C. By this time, the hyphal tips had reached a steady-state growth rate and were producing a series of tip branches without forming lateral branches. We imaged hyphal growth across the plate with transmitted light on a Zeiss Axio Zoom stereo microscope. To quantify hyphal tip growth rate over time, we measured total micrometers of growth per 30 minute intervals for individual hyphal tips. When a tip branch occurred, we continued measuring the growth rate of one of the newly formed branches. Growth rate values were measured for a minimum of 200 mins per cell. We note that there is a transient drop in tip growth rate immediately after a tip branch, however, our goal was to determine a temporally averaged growth rate, which we obtained by measuring at 30 minute intervals. We also measured spatial and temporal intervals between branching events within individual cells. As shown in the diagram in Fig. 3A of the main text, we measured branch intervals along continuous growth paths within cells, following only one of the two daughter tip branches after each branching event, since overlapping growth prevented measurement of the full distribution of branch intervals within a cell.

### Modeling Methods:

**Growing oscillator model details.** Simulations were performed using the forward Euler method with time step $\Delta t = 0.1$ minutes. Other than the example in main Figure 2B showing perfect synchrony, all model runs begin with the initial $N(0)$ oscillators given uniformly sampled random phases from $[0, 2\pi)$ at $t(0)$. Each oscillator $\theta_i$ is assigned a natural frequency $\omega_i$, randomly sampled from a Gaussian based on the experimental data as described in the main text. Each oscillator retains this natural frequency for the duration of its lifetime, and upon division at $\theta_i > 2\pi$ the daughters receive unique natural frequencies. Each run of the simulation represents a single fungal mycelial network, initialized with a different random seed. Each simulation was run for 500 minutes unless otherwise stated. The spatial positions of oscillators in main Figure 2B&C are arbitrary for the purposes of visualization.

As the initial state of our model represents a single segment of an established mycelial network (as illustrated in main Figure 2D), we add an influx to $N(t)$ to account for nuclei that would flow in from the unmodeled part of the network. Cytoplasmic flow in fungal networks occurs due to cell volume expansion at the hyphal tips and subsequent water intake throughout the network [2]. We do not know if there is spatial variability in water uptake within a cell, however we assume an influx of nuclei that is proportional to the growth rate $\alpha$, with the frequency of incoming nuclei based on the average experimental nuclear density. When nuclei enter the simulation this way they are assigned a natural frequency as described above and uniformly sampled random initial phase. They then go on to cycle and divide like the rest of the nuclear population.

**Changing model assumptions.** The model in which cell cycle progression and branching are independent of nuclear density fails to maintain constant nuclear density, even when model assumptions are altered. In Supplemental Figure 2A we show that by generating experimental natural frequency variability through noise gives the same results as main Figure 3E. Here we give each oscillator identical natural frequencies of $2\pi/120$ (min$^{-1}$), close to the experimental average. At each time step, Gaussian noise of magnitude $\sqrt{2D\,\Delta t}\,\varepsilon$ is added, where $\varepsilon \sim \mathcal{N}(0,1)$. Noise strength of $D = 0.02$ produced an average frequency distribution with a standard deviation of 40 minutes on average. In Supplemental Figure 2B we show that drawing the temporal interval between each branch based on the experimental data (rather than having new sets of branches emerge simultaneously) still fails to produce constant density.

**Adding cell cycle checkpoint.** We add a size sensing cell cycle checkpoint to the cycling of each nucleus as follows. The standard deviation of the experimental nuclear density curves was found to be approximately 1 N/100$\mu m$ (main Figure 4H). We initialize simulations to the average density value of 20 N/100$\mu m$, and choose a nuclear density threshold value based on three times the standard deviation, at 23 N/100$\mu m$. If $N(t)/L(t)$ is below the set threshold then nuclei cycle at their chosen natural frequencies. If $N(t)/L(t)$ exceeds the threshold, then any nucleus will be subject to the checkpoint if they attempt to exit G1 (defined as $[0,\pi)$). Each nucleus is then held at $\pi$ until either $N(t)/L(t)$ drops below the threshold or they stochastically escape the checkpoint. This premature escape is modeled as a Poisson process with escape rate $\lambda$, such that the probability of each oscillator exiting G1 at each time step is $\lambda\Delta t$. To avoid underestimating cycle lengths, we computed the average cell cycle length per simulation run only from nuclei born within the first 100 minutes, since late-born nuclei with longer cycle lengths may not have completed their cycle by the end of the simulation.

**Quantifying synchrony.** We quantify synchrony using the Kuramoto order parameter $r = \left|\frac{1}{N}\sum_{j=1}^{N} e^{i\theta_j}\right|$ [3]. The order parameter of all phases is calculated at individual time points and then averaged temporally to obtain a single value per simulation run. However, the expected value of the Kuramoto order parameter has a dependence on N. As our model has a dynamically changing number of oscillators, and the order parameter is sensitive to the number of phases, we calculate the order parameter from simulation time steps where $N(t) > 1000$ (Figure 2C).

**Adding density-dependent branching.** We next modify the model so that branch formation occurs in response to rising nuclear density. When $N(t)/L(t)$ exceeds the same threshold value from the cell cycle checkpoint, the next set of branches will be produced. We assume that this does not occur instantaneously, so after the density threshold is exceeded the branching event occurs at rate $\beta$, such that the probability of branch formation at each time step is $\beta\Delta t$. We also impose a minimum time interval between subsequent branch events of 70 minutes, based on the experimental distribution in main Figure 3A.

### Supplemental Figures:

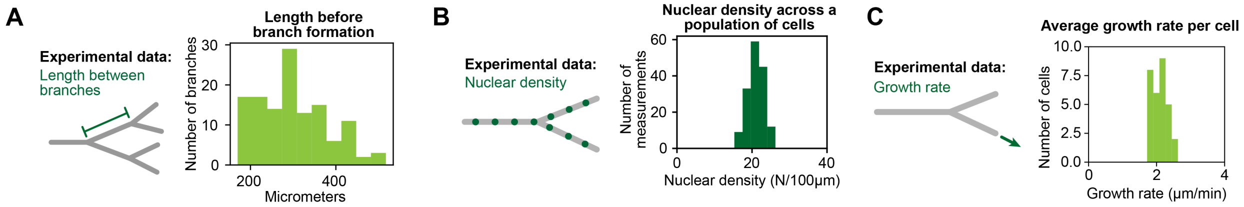


**Supplemental Fig. 1.** Experimental data for model parameters. (A) Distribution of length of hyphal segments between subsequent branch events within *Ashbya* cell networks (N = 127 hyphal segments). (B) Nuclear density across cells at single time points (N = 158 density measurements across 10 mycelial networks). (C) Average growth rate of individual hyphal tips over multiple hours of growth (N = 30 hyphae).

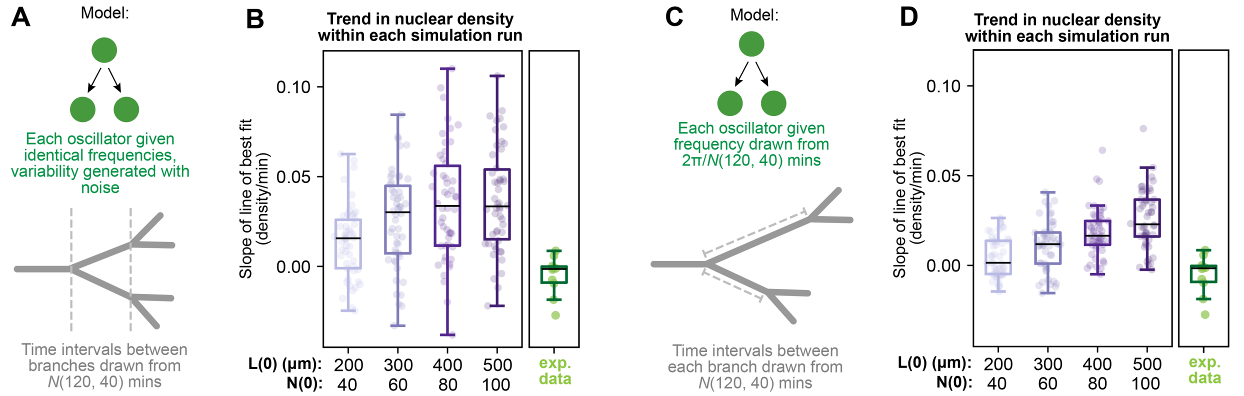


**Supplemental Fig. 2.** Matching cycle lengths and branch timing does not maintain constant nuclear density, even under different model assumptions. (A) Cycle length variability (SD=40 minutes) was generated from identical oscillators by adding Gaussian noise. Branching time intervals were drawn from a Gaussian distribution with mean and standard deviation of $120 \pm 40$ minutes. Density slopes shown for N=50 simulation runs per $L(0)$, compared to N=10 experimental mycelial networks. (B) Cell cycle lengths and time passed before each individual branch event are randomly drawn from a Gaussian with mean and standard deviation of $120 \pm 40$ minutes. N=50 simulation runs per $L(0)$, compared to N=10 experimental mycelial networks.

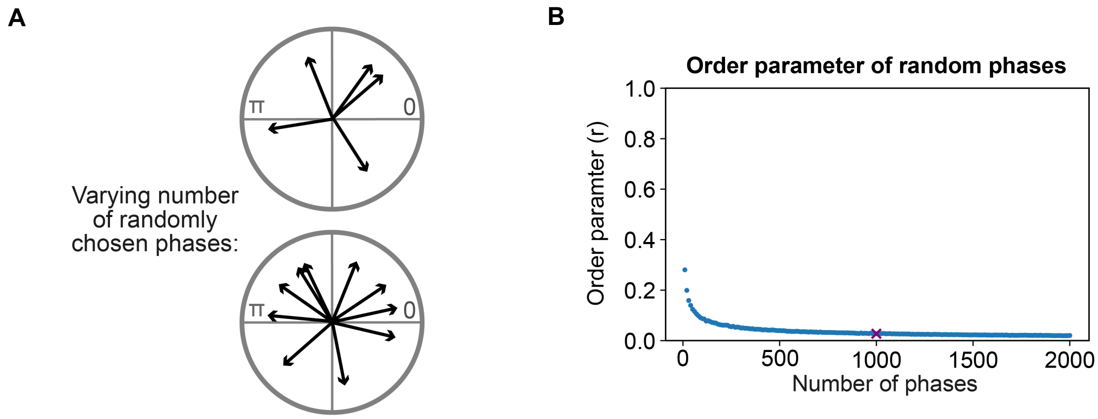


**Supplemental Fig. 3.** The expected value of the order parameter depends on the number of oscillators. (A) To illustrate this, we sampled increasing numbers of random phases drawn uniformly from $[0, 2\pi)$. (B) For each fixed number of chosen phases, we repeated this sampling 1,000 times and calculated the mean order parameter, shown here. In our model, we compute the order parameter only at time points with more than 1,000 phases.

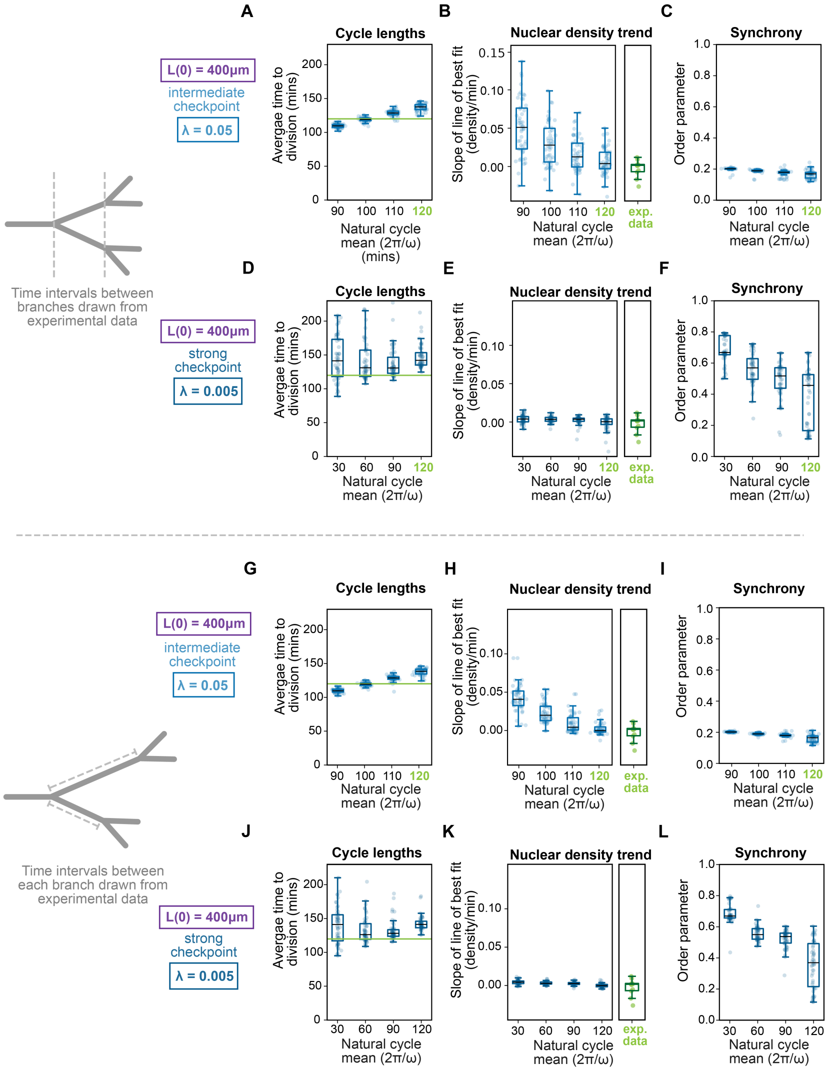


**Supplemental Fig. 4.** Effect of changing the mean natural frequency on nuclear density control and synchrony. For all simulations, $N(0) = 400$ and N=50 simulations per parameter set. (A) Showing the average cell cycle length that resulted from each simulation run. When the cell cycle checkpoint is of intermediate strength ($\lambda = 0.05$), shifting the natural frequencies leads to a linear shift in observed cell cycle lengths. Horizontal green line shows 120 minutes as reference. (B) Slopes of linear best fit to density. Decreasing cell cycle lengths worsens nuclear density control here as the checkpoint is not strong enough here to compensate. (C) Average order parameter per simulation. The systems remain asynchronous for this checkpoint strength. (D) Increasing the cell-cycle checkpoint strength ($\lambda = 0.005$) prevents the resultant cycle lengths from reaching 120 minutes, regardless of how much the natural frequencies are shifted. (E) Slopes of linear best fit to density. The strong cell cycle checkpoint prevents nuclear density from trending up. (F) Average order parameter per simulation. Synchrony arises from the strong cell-cycle checkpoint. As the cell cycle length is shortened, the checkpoint must act more frequently to reduce nuclear density, causing greater synchrony as nuclei increasingly accumulate at G1 exit. (G-L) Shows that we get similar results to (A-F) if each individual branching event occurs independently. Each simulation was run for 500 minutes, unless the cumulative number of nuclei exceeded 5000 (which occasionally occurred when cycle lengths were reduced).

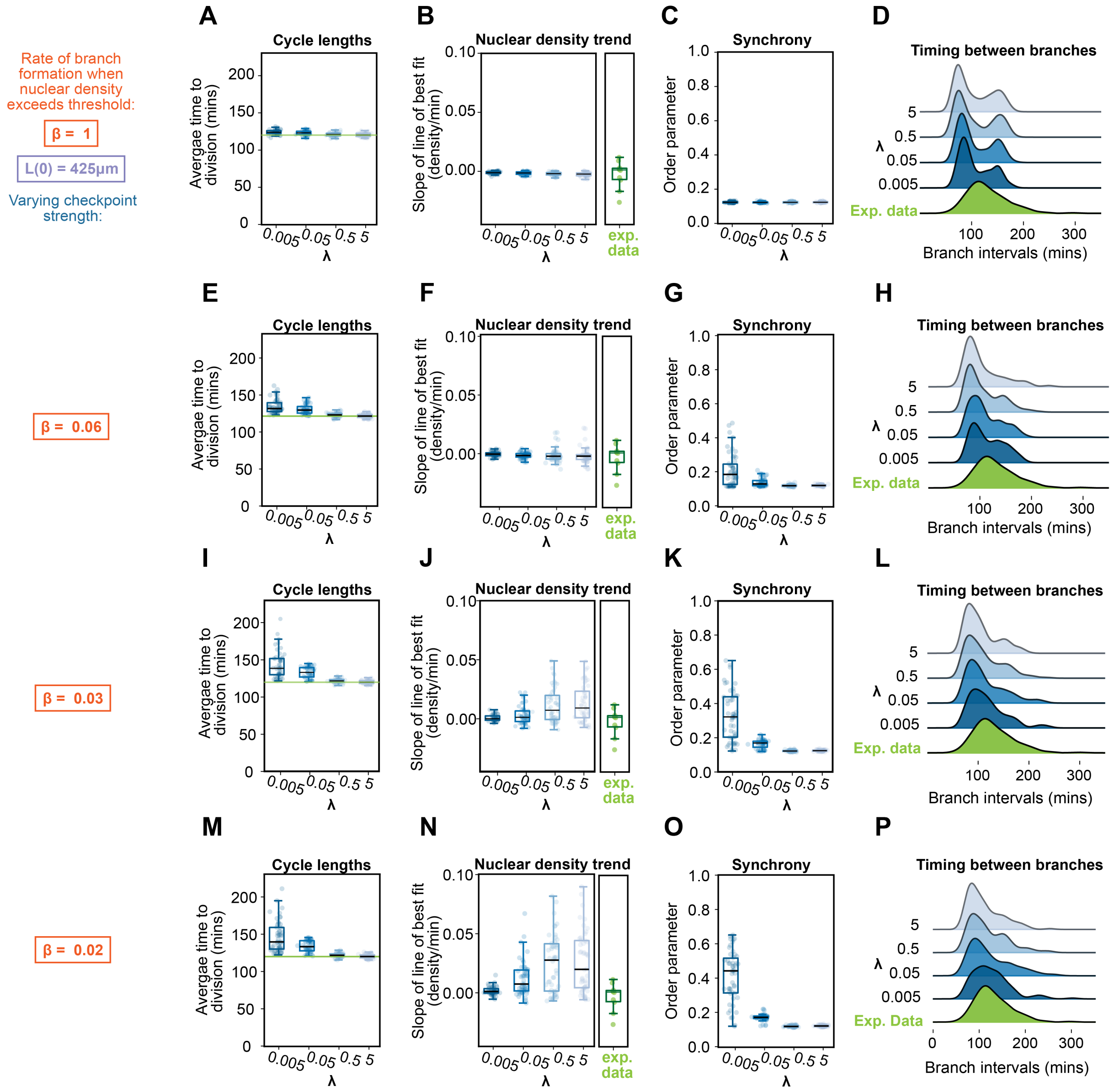


**Supplemental Fig. 5.** Trade-off between rate of branch formation and agreement with experimental data. For all simulations, $N(0) = 400$ and N=50 simulations per parameter set. (A) Rapid branch formation after density exceeds threshold ($\beta = 1$). Average resultant cell cycle length per simulation run shown. (B) When branches form quickly, density is well controlled, and (C) asynchrony is maintained. (D) Distribution of times between branch formation, however, differs from the experimental data as rapid branches form at the onset of each simulation. (E-G) Slowing average time of branch formation ($\beta = 0.06$) slightly worsens nuclear density control, (H) but begins to bring the branching time distribution closer to experimental data by shifting the distributions to the right. (I-L) $\beta = 0.03$ provides the best balance between maintaining nuclear density control and reproducing the experimental branching time distribution, specifically for intermediate $\lambda$ values. (M-P) Further delaying of branch formation worsens density control ($\beta = 0.02$ shown here).

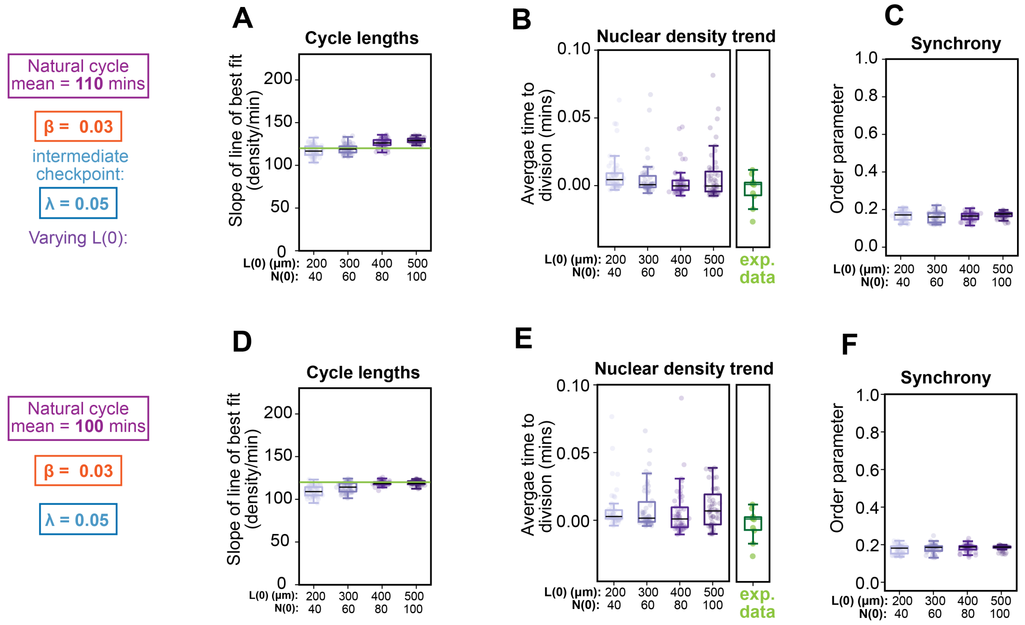


**Supplemental Fig. 6.** There is not a single shift in natural frequencies that leads to all initial conditions matching the experimental cycle lengths. For all simulations, N=50 runs per parameter set, $\beta = 0.03$, and $\lambda = 0.05$. (A) We shift the mean natural frequency to $2\pi/(110$ minutes$)$. Simulations with $L(0) = 300$ match the experimental cycle length mean, but longer initial lengths, with correspondingly higher $N(0)$ values, have longer cycle lengths. (B-C) show agreement with experimental data. (D) We next shift the mean natural frequency further to $2\pi/(100$ minutes$)$ and find that now simulations with longer $L(0)$ values match the experimental cycle length mean, (E) but have worsened nuclear density control. (F) As in C, synchrony remains low for these parameter values.
